# Unlocking the role of SH3PXD2B in epithelial-to-mesenchymal transition driving Breast Cancer Lung Metastasis

**DOI:** 10.1101/2024.10.16.618596

**Authors:** Subhajit Das, Riya Khilwani, Shailza Singh

**Affiliations:** Systems Medicine Laboratory, BRIC-National Centre for Cell Science, NCCS Complex, Ganeshkhind, SPPU Campus, Pune-411007, INDIA

**Keywords:** Breast cancer, Breast cancer lung metastasis, SH3PXD2B, systems biology, bioinformatics, therapeutics

## Abstract

Despite advancements in breast cancer treatment, metastasis remains a significant challenge, contributing to high mortality rates. This study investigates the mechanisms underlying breast cancer lung metastasis, with a particular emphasis on the role of SH3PXD2B in driving epithelial-to-mesenchymal transition. Here, we analyzed RNA sequencing data from CCLE and TCGA databases and characterized that SH3PXD2B expression is elevated in breast cancer (BC) and lung cancer (LC) tissues compared to non-transformed tissues. Bioinformatics analysis revealed that SH3PXD2B regulates several cellular processes and is associated with poor prognosis in breast cancer (BRCA) and lung adenocarcinoma (LUAD) patients. Functional experiments underscore that the upregulation of SH3PXD2B promoted BC and LC cell migration in vitro; however, its knockdown inhibited the effect. Mechanistically, the mass spectrometry analysis was utilized to pull down interacting partners of SH3PXD2B, and the findings were substantiated by immunoblotting. Additionally, our findings on immunofluorescence and gene expression analysis demonstrated the additive role of SH3PXD2B and migratory proteins in promoting breast cancer progression. Further, we performed bioluminescent IVIS imaging analysis to trace the metastatic spread of MDA-MB-468 cells and observed strong signals and a high degree of dissemination to distant organs. Collectively, our results highlight the importance of SH3 domains in aggravating breast transformation, which can therefore serve as a promising therapeutic strategy in counteracting breast-related malignancy.

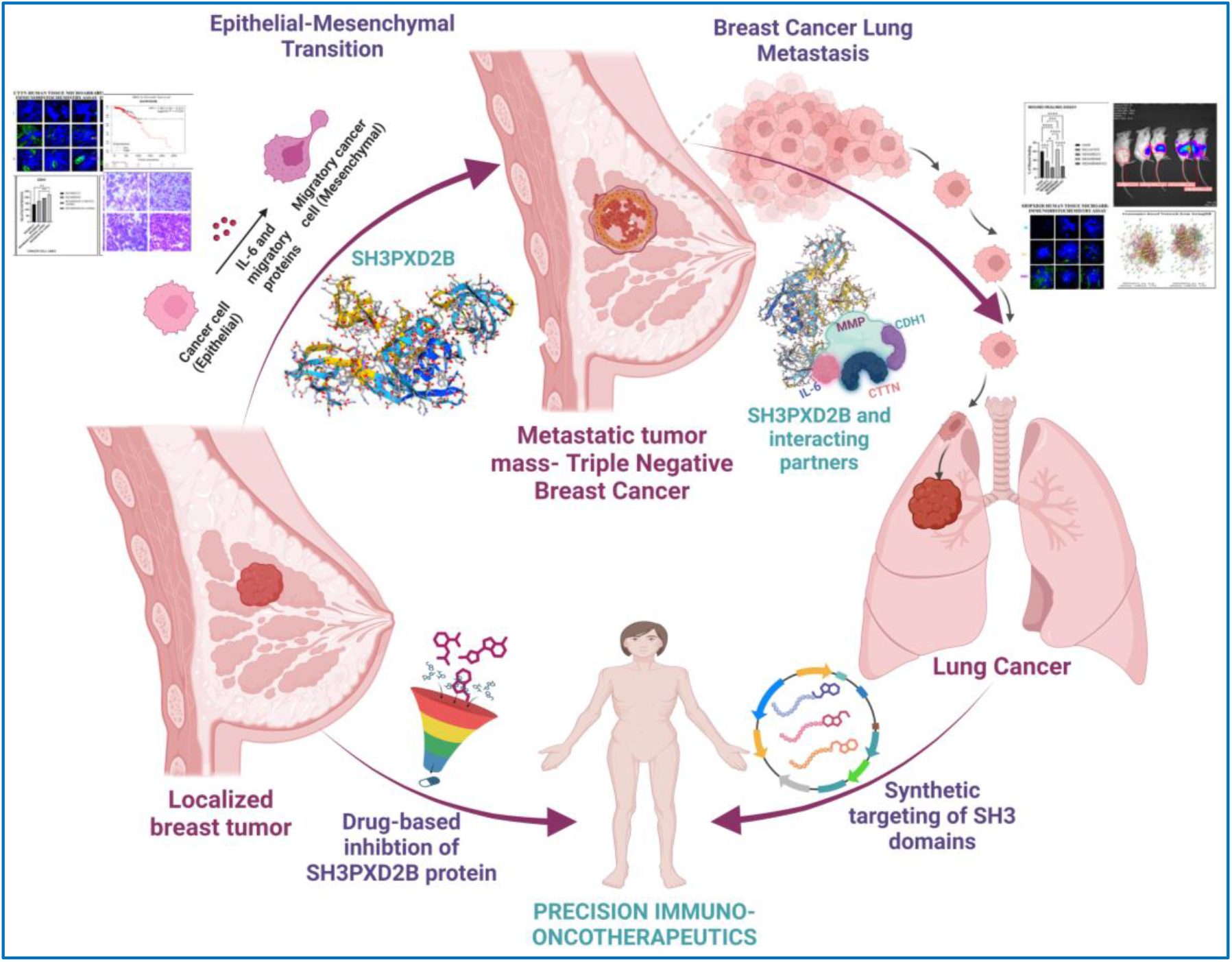

## Introduction

Breast cancer remains the most prevalent form of cancer in females and is the second leading cause of death after lung cancer. In 2022, an estimated 2.3 million breast cancer cases were reported that resulted in approximately 6,70,000 deaths (1). Of the ∼8% of the diagnosed invasive breast cancer cases, nearly 30% of patients with malignant phenotypes survive for five years (2). One-fifth of the breast cancer cases relapse, while 50-70% of the metastatic breast cancer metastasizes to distant organs, including the lung, leading to poor prognosis and therefore organ failure and death (3). Being highly heterogeneous, histological subtypes of breast cancer such as inflammatory breast cancer, invasive lobular carcinoma, medullary carcinoma, and metaplastic breast cancer are stratified by the expression of hormone receptors like estrogen receptor (ER), progesterone receptor (PR), and human epidermal growth factor receptor 2 (HER2). Each of these subtypes shows distinct bio-distribution profiling in tissues as well as inherent tumor growth tendencies (4). In lungs, the high prevalence of triple-negative breast cancer is characterized by negative ER/PR/HER2 receptors and is recognized as the most aggressive form of breast cancer contributing to high recurrence (5, 6). Despite the availability of treatment regimens for breast cancer, the prevalence and mortality related to breast cancer lung metastasis are significantly rising. This may be due to the adaptability of metastatic tumors in the lung microenvironment that alters tumoral growth dependencies. Therefore, the development of novel therapeutic strategies against metastatic breast cancer requires a comprehensive understanding of tumor-associated gene patterns that solely describe gene mediators for TNBC lung colonization and open new avenues for treatment modalities.

Metastasis is a multi-step process that requires epithelial-to-mesenchymal transition to flip immobile, closely packed epithelial phenotypes to motile, invasive mesenchymal cells with an ability to degrade extracellular matrix (7). During metastasis, the process of cellular breaching to the site of the collagen matrix is a plausible reason behind cellular invasion. This may be an effect of migrating cells expressing proteases at their leading edges important to identifying least-resistant routes while degrading extracellular matrix in breast tissues. Cellular architecture, including invadopodia, promotes matrix degeneration and stimulates the regional synthesis of protein-cleaving enzymes to cross tissue barriers (8). Multiple protein elements belonging to actin families have been discovered to modulate cell motility by coordinating the dynamics of focal adhesion and recruiting proteinases required for ECM remodeling (9). Amongst the proteins, SH3PXD2A (Tks5) is the major organizer of podosome machines that facilitate the docking of downstream adaptors for nucleation processes in plasma membrane. Recently, SH3PXD2B (Tks4), a homolog of SH3PXD2A, has been identified to play deterministic roles in influencing podosome extension and its unknown physiological effect on transformed cells. SH3PXD2B is a scaffold protein that contains four SH3 domains that facilitate protein-protein interaction, an N-terminal Phox homology (PX) domain responsible for attaching the scaffold protein via phosphoinositides to cell membranes, multiple proline-rich motifs as contact sites for SH3 domains, and the sites for Src phosphorylation (10, 11). Genetic experiments reveal the supportive role of SH3PXD2B in binding to metalloproteinases during its localization to podosomes (12). Multiple studies have shown the carcinogenic effect of SH3PXD2B in colon (13), breast (14), and melanoma models (15). These findings shed light on the explicit roles of SH3PXD2B as an invadopodial-adaptor protein associated with poor survival in cancer patients, which aids to progressive inflammation in breast-related malignancy. A recent study by Zhu et al. (2023), dictates the tumorigenic roles of SH3PXD2B that affect cancer development by forming gene regulatory networks with its interacting partners, prompting the proliferative and migratory effects of cancer cells (16). In addition to the role of SH3PXD2B, gene signatures like ADAM23, CDH1, CTTN (17), MMP2 (18), and PRPF19 (19) have been mentioned as migratory markers that serve as a pre-requisite for forming membrane ruffles and significantly alter actin dynamics by deregulating repair processes in prolonging cancer cell survival.

Herein, our research employs in silico approaches to identify SH3PXD2B expression patterns in experimental data pertaining to normal, MBC, and lung metastasis, along with the cytofunctional validation of the role of SH3PXD2B in breast cancer lung metastasis with the aim to identify potential biotherapeutic targets.

## Methods

### Data acquisition and in silico analysis

To investigate the advances in breast cancer lung metastasis, ATAC Seq and RNASeq data (GSE138122), including transcription factors, breast cancer subtypes, and their role in brain and lung cancer recurrence, were obtained from the study by Cai et al. 2020. The transcriptomics data was reanalyzed using the New-Tuxedo pipeline, Pertea et al. (2016) to identify differentially expressed genes in epithelial-mesenchymal transition. The findings were further validated by comparing them with the expression data from the Cancer Cell Line Encyclopedia and the Cancer Genome Atlas. Also, the data from the Human Protein Atlas were acquired to investigate translational signatures and binding partners of SH3PXD2B in metastatic breast and lung cancer cell lines.

### SH3PXD2B structure prediction and curation

A high-confidence protein structure of SH3PXD2B was modeled using the NCBI Accession ID: NP_001017995.1 through the AlphaFold database (Google Colab). The disordered regions in the structure were corrected by submitting a FASTA file of the SH3PXD2B protein to the Protein Disorder Prediction System. The predicted structure was analyzed to look for the protein’s secondary organizational domain using PDBsum. Further, to unravel the regulatory impact of post-translational modifications on proteins, the MusiteDeep deep learning framework was used. The results were obtained at a cutoff score of 0.04 and analyzed for post-translational modified sites on proteins.

### Gene set enrichment analysis (GSEA)

GSEA was done with the aim of identifying predefined gene sets that exhibited statistically significant differences between two biological conditions. GSEA was accessed through the Broad Institute platform to calculate an enrichment score. As recommended by the GSEA platform, a false discovery rate of 0.05 was considered a statistically significant score during analysis.

### Survival analysis

The Kaplan-Meier-plot was used to calculate the cumulative survival rate of patients over time. The breast cancer mRNA RNA-seq data in a Kaplan-Meier plotter was utilized for calculating the overall survival rates in patients. The best-performing log-rank p values, 95% confidence intervals, and hazard ratio were calculated and employed to visualize the survival curves between two cohorts.

### Transcription Factor Target Gene (TFTG) network

A bio-molecular TFTG interaction network was constructed by gathering information through literature surveys on BCLM-specific proteins and transcription factors. The related transcription factors were obtained from the TF2DNA and TF-link databases. Cytoscape (v.3.4.0) was used to construct the network, followed by analysis using various algorithms in CytoHubba.

### IP/MS proteomics analyses

The cultured MDA-MB-231 and MDA-MB-468 cells were washed and resuspended in NETN lysis buffer containing 50 mM Tris (pH8), 250 mM NaCl, 5 mM EDTA, and 0.5% NP-40 with phosphatase and protease inhibitors. Following 20 min incubation, the samples were vigorously vortexed and, centrifuged at 16,000 g for 20 min at 10 °C and the protein fraction was collected for BCA estimation. Precisely, the impurities in cell lysates were removed by incubating samples at 4 °C for 1 h in 25 μl of BioRad SureBeads. Next, 2 μg of SH3PXD2B antibody and rabbit polyclonal IgG (-ve control) were added to 500 μl of PBST consisting of 50 μl of cleaned BioRad Protein G magnetic beads. The samples were kept in continuous stirring conditions for 4 h and then rinsed multiple times to remove the unbound fraction. Additionally, the cells were washed with 500 μl of 100 mM ammonium bicarbonate buffer and subjected to the elution process. Briefly, the process involves digesting protein complexes using 0.1% RapiGest and 5 mM DTT for 21 h, followed by the cleaning step, which involves filtering protein through a 3KDa column. The filtered extract was further digested with 1 μg of trypsin for 18 h at 37 °C, and the tryptic peptides were analyzed after they underwent desalting procedures. The proteins in a resultant sample were acquired using mass spectrometry employing the Thermo Scientific™ Orbitrap fusion Tribrid LC-MS/MS system. While analysis using the Proteome Discoverer 2.2 tool, strict statistical standards was maintained for unique peptides and protein FDR confidence levels.

### Pathway enrichment analysis

The pathway enrichment study for metastatic markers in breast cancer was conducted using the Cytoscape plugin-Biological Networks Gene Ontology tool (BiNGO), which computed the enrichment score of biological markers or gene ontologies using the transcription factor-target gene network. A hypergeometric test was used to calculate the p-value, and the obtained false positive values were corrected using the Benjamin Hochberg test. Additionally, functional enrichment analysis was achieved using StringDB and Webgestalt to validate experimental results.

### Cell culture and maintenance

Human highly metastatic breast cancer cell line (pleural origin), MDA-MB-468; human breast cancer mesenchymal cell line, MDA-MB-231; human epithelial breast cancer cell line, MCF-7; and human lung adenocarcinoma cell line, A549, were procured from the cell repository NCCS (National Centre for Cell Science), Pune. Human breast non-tumorigenic epithelial cells, MCF-10A (CRL-10317); human lung epithelial cell line, Beas-2B (CRL-9609); and human lung adenocarcinoma cell line, H1975 (CRL-5908), were procured from the American Type Culture Collection (ATCC). The MDA-MB-231 and MDA-MB-468 were cultured in L-15 medium supplemented with 15% FBS (Gibco), 100 U/ml of penicillin, and 100 μg/ml of streptomycin. MCF-7 cells were cultured in DMEM containing 10% FBS and the antibiotic combination described above. MCF-10A cells were maintained in DMEM/F12 medium containing horse serum, EGF, cholera toxin, insulin, and penicillin-streptomycin. H1975 and Beas-2B cells were cultured in RPMI medium containing 10% FBS, 100 U/ml of penicillin, and 100 μg/ml of streptomycin. A549 cells were cultivated in Ham’s F-12 medium, 10% FBS, 100 U/ml of penicillin, and 100 μg/ml of streptomycin. Culture conditions for all the cells were maintained at 37 °C in 5% CO_2_, except for MDA-MB-231 and MDA-MB-468 cells, which were maintained without CO_2_ supplementation. After the cells reached 80-90% confluency, they were dissociated using TPVG and centrifuged at 300 g for 5 min for further processing.

### Protein extraction and western blotting

Briefly, total cellular protein was harvested and lysed in RIPA buffer containing 10 mM tris base (pH 7.4), 7 M urea, 2 M thiourea, and 2% sodium deoxycholate and quantified using the Pierce™ Bradford Plus Protein Assay (# 23228). Potein lysates were equally separated on SDS-PAGE and transferred to PVDF membranes (Hi-FiBloE™ PVDF Membrane for Blotting). The membranes were then blocked with 5% non-fat skim milk and incubated with primary antibodies. HRP-conjugated secondary antibody was added, and membranes were visualized using an ECL detection reagent (Novex™ ECL Chemiluminescent Substrate Reagent Kit, WP20005). The quantity of protein markers was analyzed through Cytiva Amersham ImageQuant 500 equipment. The protein expression level was normalized to the loading control, β-actin.

### Antibodies and reagents

Antibodies for SH3PXD2B (PA5-57673, 1:1000), MMP14 (MA5-38574, 1:1000), PRPF19 (15414-AP, 1:1000), and SNAI1 (PA5-23482, 1:1000) were from Thermo Fisher Scientific (Waltham, Massachusetts, US). CTTN (h222-3503, 1:1000) and β-actin (4967, 1:1000) antibodies were from Cell Signaling Technology (Boston, MA, USA). Antibodies for MMP2 (A6247, 1:1000) and ADAM23 (A14263, 1:1000) were purchased from ABclonal (Cummings Park, United States). CDH-1 (sc-7870, 1:1000) was obtained from Santa Cruz Biotechnology (Dallas, Texas, United States). The respective isotype controls for antibodies were procured from Cell Signaling Technology (Boston, MA, USA).

### Cell migration assay

Wound healing and a Boyden or Transwell chamber assay were used to test the migratory ability of control vs. knockdown cancer cells. In a wound healing assay, a gap in a confluent monolayer of transformed lung epithelial cells was created to replicate a wound. Briefly, 1*10^5^ cells were seeded in a 24-well plate until monolayer formation, and a wound was made using a 1 mm-wide tip. Micrographs were taken at 0 and 24 h intervals (x10) using an Olympus FV3000 scanning laser microscope and quantified using image analysis software, ImageJ. The transwell assay was performed in a specialized chamber separated by a porous membrane (CLS3464-48EA, Sigma-Aldrich). The cells were seeded in the upper chamber, and a chemoattractant serum was placed in the bottom chamber for 24 h to assess the cell’s stimulating effect. After incubation, the cells were stained with a 0.4% trypan blue solution (15250061, Gibco) and counted under a Nikon inverted microscope.

### Plasmids

The expression plasmid pLentipuro3/TO/V5-GW/EGFP-Firefly Luciferase encoding EGFP was obtained from Addgene (Watertown, MA, USA). The shRNA constructs specifically targeting SH3PXD2B were analyzed in the TRC library database of the RNAi Consortium shRNA library (Broad Institute, Cambridge). After analysis, the plasmid pLKO.1 encoding SH3PXD2B shRNA was obtained from the Indian Institute of Science, Bangalore. The provided scrambled shRNA control construct was referred to as a negative control for experimental sets.

### shRNA knockdown

Short hairpins targeting human SH3PXD2B and scrambled control (shScr) were used in the study to generate SH3PXD2B knockdown in the MDA-MB-468 (Breast Cancer Lung Metastatic) cell line. Briefly, liopfectamine 3000 was used to introduce shRNA constructs into breast cancer cells. During the transfection process, the ratio of shRNA to liposomes and the incubation time were systematically evaluated to determine optimal transfection conditions. Following transfection, successful delivery of the constructs was confirmed using immunoblotting. To note, of all the analyzed shRNA constructs, SH3PXD2B shRNA with clone ID ‘TRCN0000147775’ was selected to knockdown the expression of cellular SH3PXD2B.

### Cell viability assay

In a study, a cell viability assay was performed to test the effect of the PI3-Kinase (IC_50_ = 5 nM) inhibitor, Wortmannin (CAS 19545-26-7, Sigma-Aldrich). Briefly, after treatment of cultured cells with Wortmannin, they were incubated with 3-(4,5-dimethylthiazol-2-yl)-2,5-diphenyltetrazolium bromide (MTT) for 4 h, and the optical density was measured using a microplate reader to detect formazan crystal formation.

### Tissue Microarray

An in vivo tissue microarray study was performed on a Human Breast Tissue MicroArray (Cancer), NBP2-30212, that was procured from Novus Biologicals USA. The samples were pre-arrayed on a slide, having a thickness of 4 μm with a core diameter of 2.0 mm, and were ready to be used for immunohistochemistry analysis. The panel comprises 59 samples, including 40 samples with infiltrating duct carcinoma, 10 samples with metastatic carcinoma, and 9 samples representing normal healthy subjects (NHS). The tissue was labeled against ADAM23, CTTN, IL-6, MMP2, and SH3PXD2B antibodies to assess their metastatic potential. The arrayed samples from healthy individuals served as a reference for the experimental set.

### Mice model

NOD-SCID mice, 6-8-week-old females, were reared and bred in the experimental animal facility (Project No. IAEC/2023/B-452) of the National Centre for Cell Science, Pune, India. The use of animals was followed as per the protocol approved by the Institutional Animal Ethics Committee (NCCS) registered under CCSEA.

### Bioluminescence luciferase assay

An in vitro functional luciferin protocol was adopted to evaluate the bioluminescent properties of stably transfected cell lines. Briefly, HEK293 FT cells were cultured and underwent the general PEI transfection protocol, and the resultant viral supernatant was collected post 48 h and 72 h of transfection and filtered using 0.45 μm filter unit. The supernatant was added to the grown MDAM-MB-231 and MDA-MB-468 cells and allowed to rest for 24 h. After incubation, the spent medium was exchanged with a fresh medium, and puromycin (2 μg /ml) was added to generate stable mammalian cell lines, MDA-MB-231luc2 and MDA-MB-468luc2, respectively. After 2-3 passages, the cells were treated with 150 μg/mL of D-luciferin substrate (IVISbrite D-Luciferin Potassium Salt Bioluminescent Substrate, XenoLight, Revvity), and the luciferase expression was measured using an Olympus FV3000 confocal laser scanning microscope.

### In vivo biodistribution analysis of BCLM

For the protocol, disease-free NOD-SCID mice were injected with 2*10^6^ MDA-MB-231luc2 or MDA-MB-468luc2 cells through two routes. Of which, 2*10^6^ cells were injected through the tail vein route for widespread distribution and 1.8*10^6^ cells were injected through the mammary fat pad for localized tumor formation. The injected cells were allowed to grow, and their cellular metastatic ability was evaluated. After 7 days, the mice were administered intraperitoneally with a dose of 150 mg/kg of D-luciferin substrate and subjected to imaging while under anesthesia. The bioluminescent imaging was performed utilizing the PerkinElmer IVIS® Spectrum 2 optical imaging platform, and the analysis was conducted using the Living Image program v4.8.2 (PerkinElmer).

### Statistical analysis

GraphPad Prism 10.0.1 was used to perform statistical analysis. The data are represented as mean ± SEM, and each data set is the representative outcome of at least three independent experiments. Accordingly, two-tailed unpaired Student’s *t* tests, Welch’s t-test or one-way or two-way ANOVA with Tukey’s correction or anova multiple comparisons were used. *, *p* < 0.05; **, p<0.01; ***, p<0.001; ****, p<0.0001 were considered significant.

## Results

### SH3PXD2B drives EMT by aiding cell migration

A previous study (20) has revealed that the chromatin landscape of HER2-enriched breast cancer subtypes activates specific transcription factors that modulate the epigenetic states responsible for metastatic colonization in the brain and lung. In the study, to investigate the processes involved in BCLM, we first reanalyzed the additional transcriptomics data (GSEA138122) using the New-Tuxedo pipeline, and the data revealed a higher expression of SH3PXD2B than other proteins in BCLM (Figure 1A). To validate this finding, the expression data from the CCLE and TCGA databases was comparatively investigated. As shown in figure 1B, higher expression of SH3PXD2B was detected in the highly metastatic breast cancer cell line, MDA-MB-468, over the TNBC-metastatic cell line, MDA-MB-231, followed by its expression in lung adenocarcinoma cells, which correlates the role of SH3PXD2B in metastasis. Additionally, analyzing the translational signatures demonstrated the higher expression of essential migratory markers like E-cadherin (CDH1), β-catenin (CTTNB1), snail (SNAI1), and slug (SNAI2) in both breast and lung cancer patients. Of which, SNAI1 and CTTN were the highest among all in BC and LC patients, respectively. Moreover, the data reveals the localization of SH3PXD2B in the breast glandular and myoepithelial cells and cellular interactions that facilitate cancer progression. However, in lung cancer, alveolar cells and myoepithelial cells are specific to SH3PXD2B function (Figure 1C). These results indicate the tissue-specific effects of co-partnering of SH3PXD2B, which is expressed at greater levels in BCLM and influences the EMT process.

**Figure 1:**
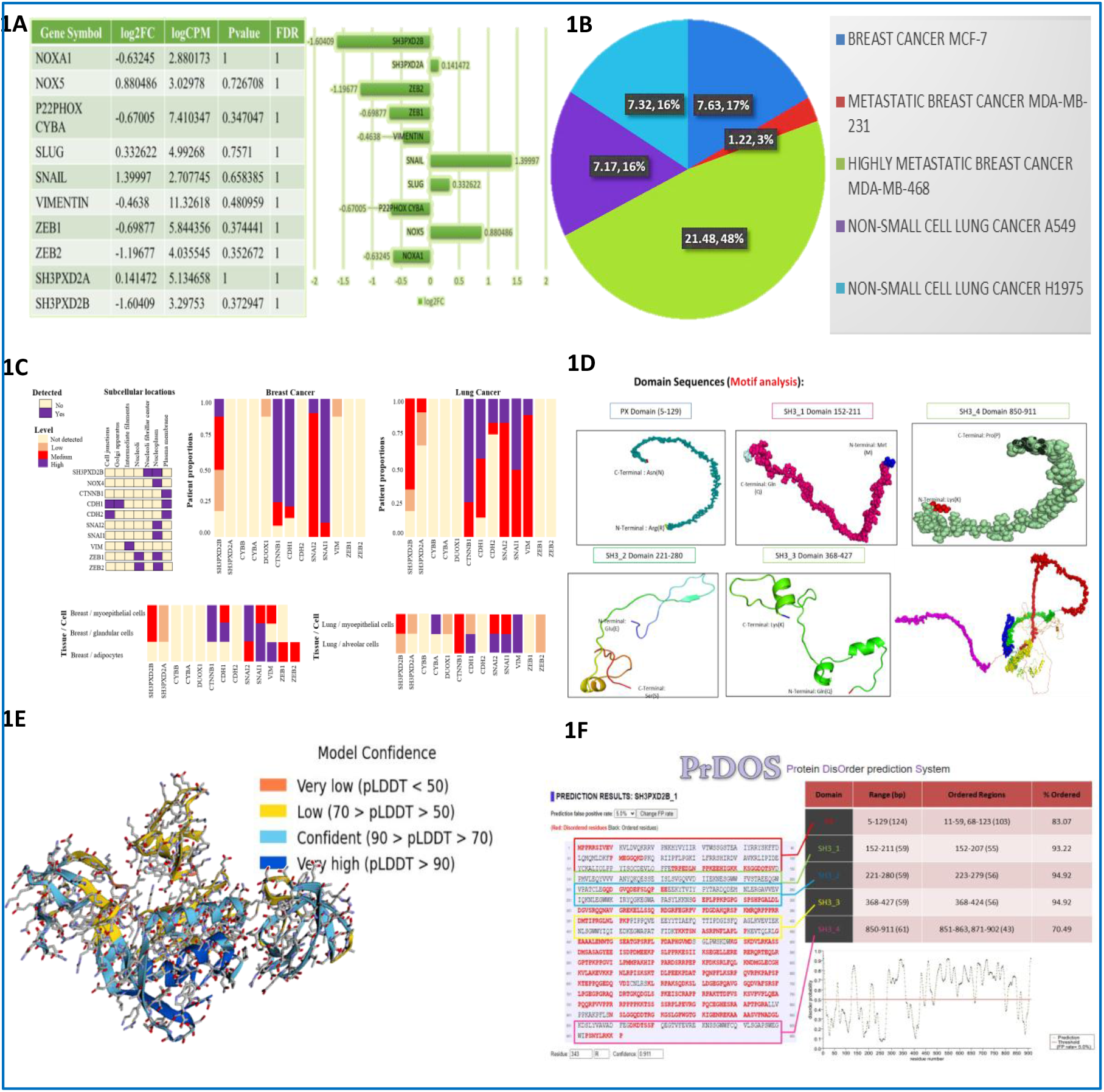
Interactions associated between SH3PXD2B and other proteins drive malignant transformation in breast and lung cancer cells. (1A) illustrates the enrichment of SH3PXD2B and other EMT genes in BCLM. (1B) A pie diagram depicts SH3PXD2B mRNA expression in different BC and NSCLC cells. (1C) The data on protein expression describing the localization of SH3PXD2B within the BC and LC cells. (1D) SH3PXD2B motif identification using MEME Suite (1E) AlphaFold2-derived structure of SH3PXD2B (1F) Prediction of disordered regions in the derived structure using PrDOS.

Since our observation that SH3PXD2B regulates the BCLM process, we used a homology modeling approach to predict the protein structure (NP_001017995.1); however, the resultant protein had low coverage. Therefore, we used the MEME suite as a bottom-up method to identify motifs (Figure 1D), and the outcomes demonstrated the presence of one PX and three SH3 domains. We then submitted these domains in an AlphaFold2 using Google Colab to achieve model confidence. The yellow-labeled regions in the ribbon and stick model relate to interfaces in SH3PXD2B that show low confidence; notably, the fourth SH3 domain is one such interaction. However, the blue section reflects higher confidence, where the pLDDT score ranges between 70 and 90 and even higher in some cases. This includes both the SH3_2 and SH3_3 domains (Figure 1E and supplementary data 1a), suggesting their functional role in metastasis. According to PrDOS estimates, the domain conservedness was indicated to be relatively high; however, the projected probability score was calculated for every disordered region in the structure. Analyzing Bokeh plots showed that SH3_2 and SH3_3 domains had the least disordered probability with nearly 95% ordered regions (Figure 1F), which also corroborated our findings of model confidence. Collectively, these results demonstrate that the functionality of SH3PXD2B relies on SH3_2 and SH3_3 domains, which might be responsible for promoting the interaction of SH3PXD2B with cell migratory proteins during BCLM.

### SH3PXD2B’s co-partnership with metastatic proteins potentiates pulmonary transformative events, resulting in decreased longevity

Given the exceptional recognition of SH3PXD2B in breast cancer, we used GSEA to predict its differential expression in both normal and malignant breast cells, leveraging the CCLE expression database. In the study, the gene enrichment score was calculated for three gene set combinations: MCF-7 (metastatic), MDA-MB-231 (TNBC), and MDA-MB-468 (TNBC pleural) with MCF-10A. Within the cancer subtypes, SH3PXD2B is negatively enriched in MCF-10A (non-cancerous condition) and decreases rapidly with an enrichment score of -0.8, -0.8, and -0.7 for the above-mentioned combinations, suggesting a higher likelihood of the involvement of SH3PXD2B in breast-related malignancy. Additionally, the butterfly plot also shows the inverse correlation of the negatively ranked genes associated with aberrant SH3PXD2B expression. The heat map shows the higher expression of SH3PXD2B gene targets for MCF-7A, MDA-MB-231, and MDA-MB-468, correlating its role in breast cancer progression (Figure 2A). Based on the role of SH3PXD2B in BCLM and its gene expression signature found in breast cancer cells, we hypothesized that the co-partnering of SH3PXD2B with its gene target might influence the process of breast cancer lung metastasis. We assigned different gene signatures (SH3PXD2B, SH3PXD2A, MMP9, CTTN, NOX5, EGF, SRC, GRB2, ZEB1, ZEB2, DUOX1, CDKN2A, SNAI1, ADAM15, MMP14, and ADAM12) to a breast/lung cancer mRNA seq database for a Kaplan-Meier survival analysis (KMplot.com). Remarkably, we found that for every gene, log-rank P value was less than 0.05, suggesting its significance in metastatic events and the occurrence of death within the allotted time frames. The worst overall survival was seen in breast cancer cases with highly expressible transcription factors, with a log-rank p-value five times lower than 0.05. Similarly, in lung cancer, patients with higher expression levels of metastatic proteins with a significantly lowered p-value demonstrated the lowest overall survival. Genes like MMPs, ADAMs, SH3PXD2A, and SH3PXD2B are crucial elements of the metastatic niche and therefore might serve a role in driving the spread of tumor cells from breast to lung (Figure 2B and supplementary data 2a). According to z-scores, genes including SH3PXD2B and DUOX2 mark the early stage of breast and lung cancer adenocarcinoma (BRCA_F1 and LUAD_F1, F1 represents Female 1). Genes like SH3PXD2A, MMP9, ZEB1, ZEB2, TP53, and CDKN2A (BRCA_F2 and LUAD_F2) are crucial in advancing the process of BCLM. However, molecular signatures involving MMP9 and GRB2 (BRCA_F3 and LUAD_F3) served as critical to progressing tumorigenic events during later stages (supplementary data 2b and 2c).

**Figure 2:**
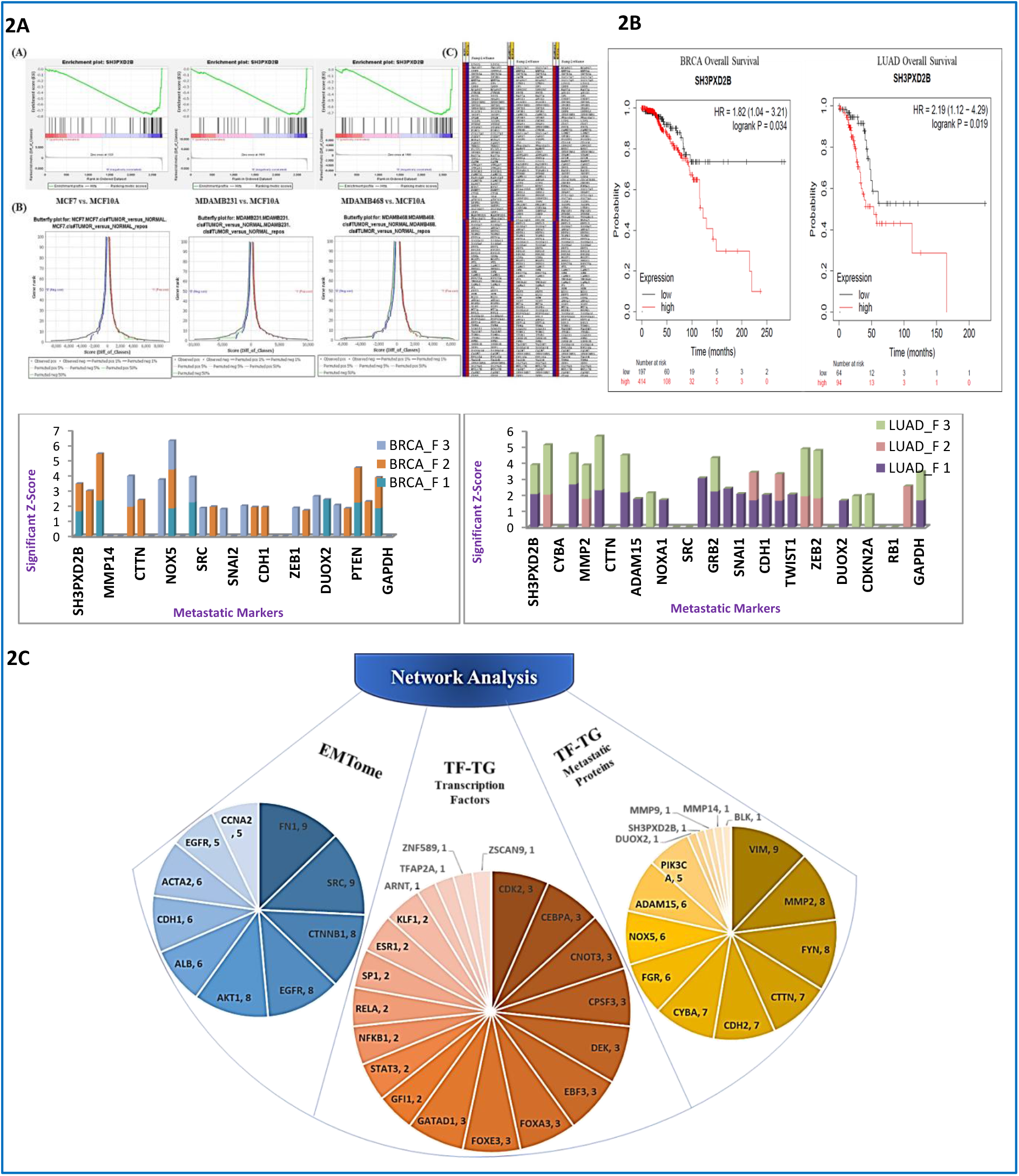
Differential expression analysis reveals SH3PXD2B complexes with other proteins during breast transformation. (2A) A figure illustrates the gene-set analysis, including enrichment plots, butterfly plots, and heat maps for all three combinations, MCF7 vs. MCF10A, MDAMB231 vs. MCF10A, and MDAMB468 vs. MCF10A. (2B) Overall survival analysis of SH3PXD2B in BRCA and LUAD and the associated z-scores for different metastatic markers involved (2C) TFTG and EMTome network analysis of BCLM markers.

To delineate the complex regulatory mechanisms underlying EMT in BCLM, we performed EMTome network and TFTG network analysis. In brief, EMTome network analysis helps to visualize and understand the interactions between core EMT genes, transcription factors, and associated signaling pathways. However, the TFTG network focuses on the interactions between TFs and their target genes, which are important in gene regulation. Here, the GSE138122 DESeq data and TFTG network-derived transcription factors and metastatic proteins were analyzed using the Cytoscape plugin, CytoHubba. Based on twelve different scoring methods, including Betweenness, Bottleneck, Closeness, Clustering Coefficient, Degree, Density of Maximum Neighbourhood Component (DMNC), EcCentricity, Edge Percolating Coefficient (EPC), Maximal Clique Centraity (MCC), Maximum Neighborhood Component (MNC), Radiality, and Stress, the top ten TFs/TGs were determined according to their frequency of occurrence. The analysis showed the enrichment of metastatic proteins, VIM, MMP2, FYN, CTTN, CDH2, CYBA, FGR, NOX5, ADAM15, PIK3CA, DUOX2, SH3PXD2B, MMP9, MMP14, and BLK; and transcription factors, CDK2, CEBPA, CNOT3, CPSF3, DEK, EBF3, FOXA3, FOXE3, GATAD1, GFI1, STAT3, NFKB1, RELA, SP1, ESR1, KLF1, ARNT, TFAP2A, ZNF589, and ZSCAN9 in the entire network. Moreover, the EMTome network analysis revealed those EMT markers, FN1, SRC, CTNNB1, EGFR, AKT1, ALB, CDH1, ACTA2, EGF, and CCNA2, which are involved in the process of invadopodia formation during metastasis (Figure 2C and supplementary data 2d). Altogether, these findings suggest the potential role of SH3PXD2B in metastasis, where it co-partners with different transcription factors and facilitates the process of BCLM, resulting in a poor survival rate of patients with both BC and LC.

### Reduction in SH3PXD2B levels upregulates translation signatures in BCLM

Our functional experiments revealed that SH3PXD2B plays a role in regulating several cellular processes that can lead to cellular transformation. Based on these findings, we hypothesized that SH3PXD2B’s interaction with other proteins might influence its metastatic potential.

To address this question, we attempted to investigate the binding partner of SH3PXD2B from breast cancer cells. CO-IP was used to identify direct interactors of SH3PXD2B in MDA-MB-231 and MDA-MB-468 cells, and translational signatures were validated by immunoblotting. Mass spectrometry analysis revealed that SH3PXD2B not only has higher expression levels in highly metastatic breast cancer cells but also has more binding partners. Of the total identified 249 interacting proteins, 106 proteins were found in the MDA-MB-468 cells. However, 64 protein hits were obtained for MDA-MB-231 cells, and 79 proteins were common in both groups (Figure 3A and supplementary data 3a). Next, we utilized StringDB to construct SH3PXD2B-protein interaction networks, and the clustering coefficient signifies that the networks are sparsely connected (Figure 3B). However, the gene ontology plot reveals that despite being critical in regulating tumor development, SH3PXD2B also regulates the processes of cell communication, development, motility, and differentiation (Figure 3C).

**Figure 3:**
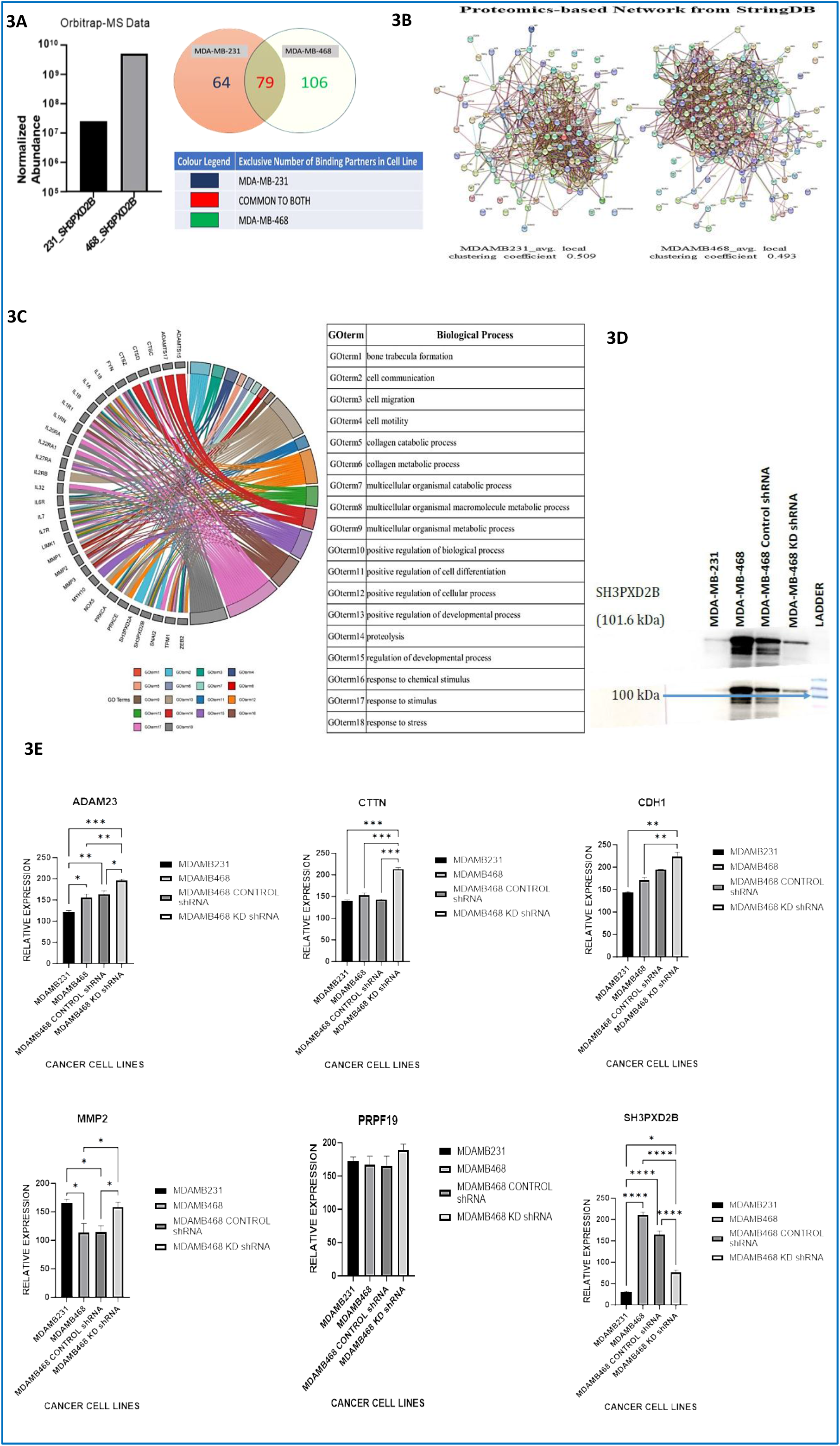
SH3PXD2B knockdown and its effect on related translational signatures: (3A) Venn diagram showing the number of common and exclusive binding partners in respective cell lines. (3B) StringDB-based network construction of SH3PXD2B binding partners (3C) Gene Ontology Circus plot shows the enriched biological pathways from the IP-MS proteomics study. (3D) Immunoblot analysis illustrates the expression of SH3PXD2B in different experimental conditions (3E) Densitometry analysis of ADAM23, CTTN, CDH1, MMP2, and PRPF19 upon SH3PXD2B knockdown.

Further, to study the roles of SH3PXD2B in breast cancer and its effect on protein partners, we knocked down its expression separately in MDA-MB-468 cells and detected knockdown efficiency using western blotting (Figure 3D). Meanwhile, the expression levels of SH3PXD2B binding proteins were checked. Immunoblotting analysis reveals that the expression levels of ADAM23, CTTN, CDH1, and MMP2 were significantly upregulated upon SH3PXD2B knockdown, meaning that these proteins being SH3PXD2B’s binding partners compensate for its loss during malignant states. Despite the changes in above-mentioned protein expression, we did not observe any significant change in PRPF-19, a component involved in the synthesis of mature mRNA (Figure 3E). These findings substantiate our previous findings on the interactions between SH3PXD2B and proteins influencing metastasis, suggesting their additive role in promoting breast cancer lung metastasis.

### SH3PXD2B knockdown in MDA-MB-468 cells inhibits breast cancer cell migration

To further investigate the cancer cell’s migratory behavior and corroborate earlier findings on their metastatic potential, we performed transwell migration and scratch wound experiments. Consistent with the findings, the transwell migration assay showed a higher migration rate for MDA-MB-468 cells over MDA-MB-231, which makes these cells a key determinant during metastatic events (Figure 4A).

**Figure 4:**
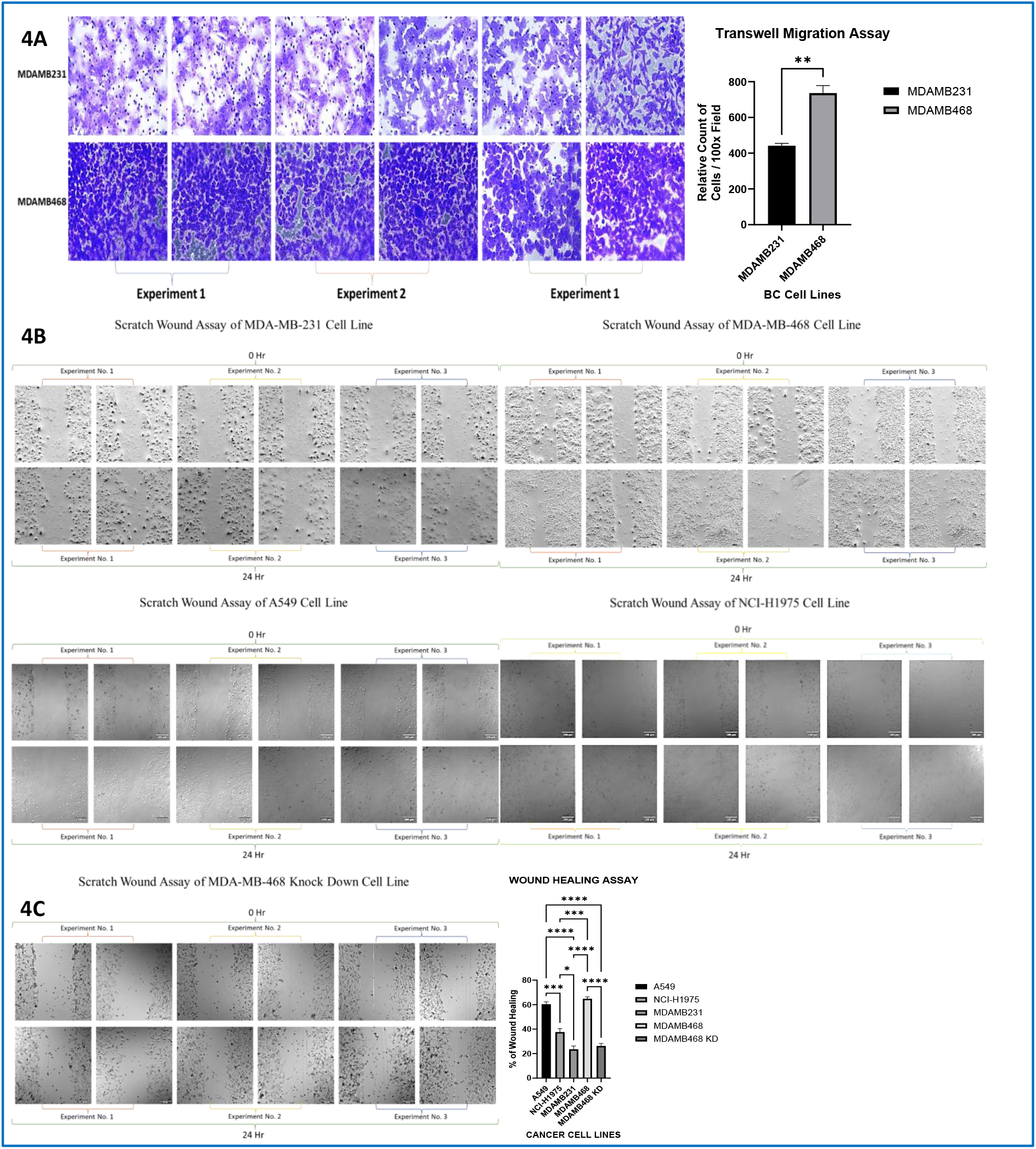
SH3PXD2B promotes BC cell migration. (4A) Quantitation of the migratory abilities of metastatic breast cancer cells (4B) Scratch wound healing effect on the migration of BC and LC cells. (4C) Quantitative analysis of the effect of SH3PXD2B knockdown on BC cell migration.

Since our observation that MDA-MB-468 is highly metastatic and is a critical driver during BCLM, we assessed the migration ability for both breast cancer cells, MDA-MB-231 and MDA-MB-468, and lung cancer cells, A549 and H1975. In line with the findings, breast and lung cancer-associated MDA-MB-468 and A549 cells, respectively, exhibited significant migration (Figure 4B). Since SH3PXD2B has been found to influence the BCLM process, we speculated that it might be a crucial player behind progressive transformation. Therefore, we assessed the migration ability of cells upon SH3PXD2B knockdown. Interestingly, we found that the knockdown of SH3PXD2B in MDA-MB-468 cells markedly inhibited the migratory abilities of breast cancer cells, resulting in greater wound width (Figure 4C, supplementary data 4a, and 4b). The resistance of MDA-MB-468 cells to SH3PXD2B knockdown, coupled with its observed effect on cell migration, suggests that SH3PXD2B plays a crucial role in potentiating transformative and metastatic events in these cells.

Given the critical role of the PI3K/Akt pathway in breast cancer progression, inhibiting it with wortmannin has been explored clinically. Herein, we employed cell viability assays and checked the dose-dependent inhibition of wortmannin on breast cancer cells. Our cell viability assay demonstrated that wortmannin, at a concentration of 400 nM, significantly reduced the viability of breast cancer cells (supplementary data 4c), suggesting its potential as a therapeutic agent. Therefore, in vitro results indicate that a combined approach of shRNA-mediated gene silencing and pharmacological protein targeting holds promise for reversing breast cancer-related malignancy.

### Immunofluorescence and gene expression data validate SH3PXD2B as a marker for breast transformation

To understand the mechanism underlying the metastatic effect of SH3PXD2B and the few TFTG-derived proteins in breast cancer, we performed immunofluorescence analysis in a breast cancer microarray panel (procured from Novus Biologicals). Normal cells exhibited negligible levels of SH3PXD2B, IL-6, CTTN, and MMP2, while breast cancer and metastatic breast tissues showed significantly higher expression levels. In contrast, ADAM23 had the highest expression in non-transformed cells, which gets reversed in malignant states, suggesting its potential role as a tumor suppressor (Figure 5A).

**Figure 5:**
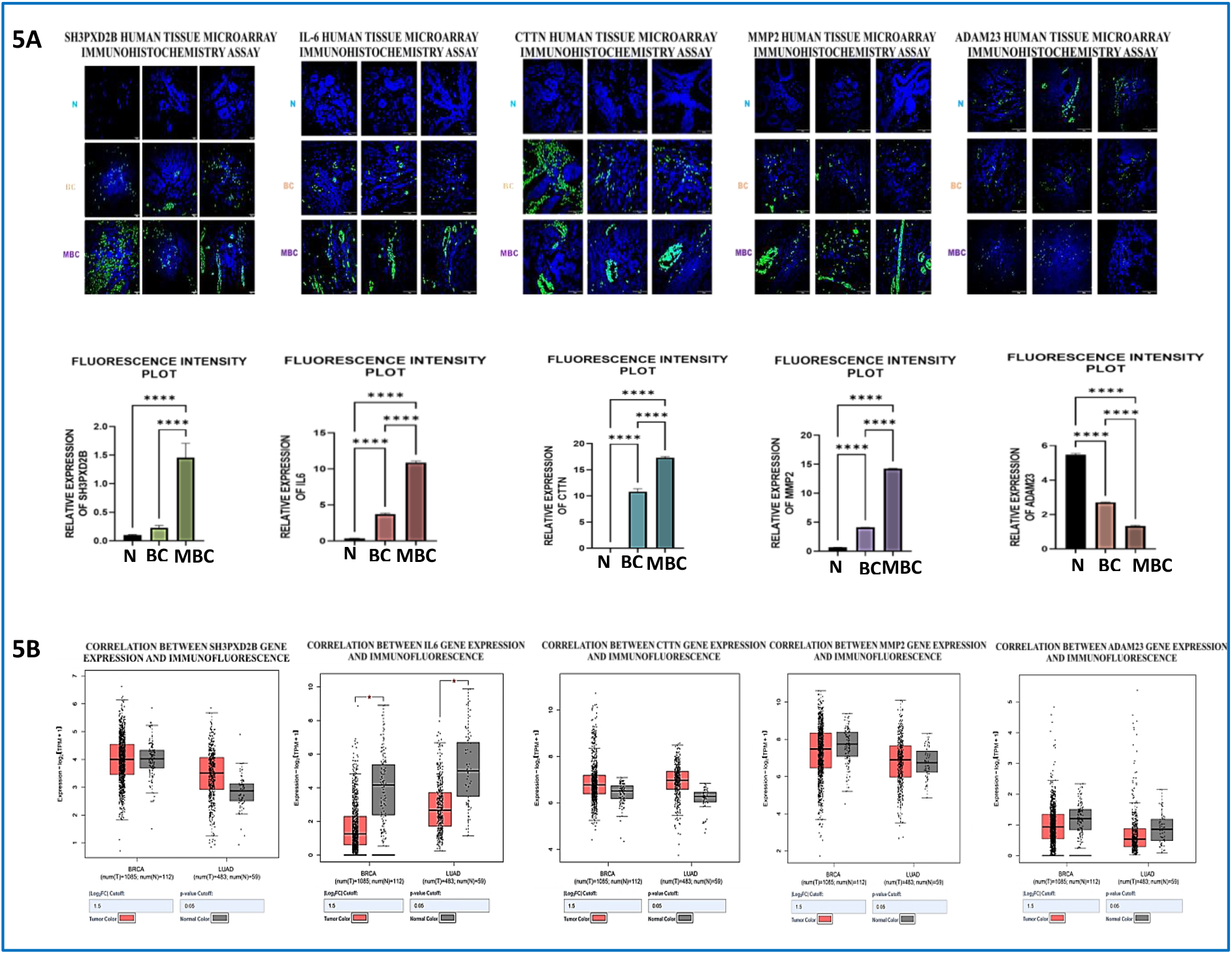
Validation of SH3PXD2B and other metastatic markers using immunofluorescence and gene expression analysis. (5A) illustrates the expression profiles of SH3PXD2B, IL-6, CTTN, MMP2, and ADAM23, respectively. While (5B) represents the gene expression analysis of respective markers.

Furthermore, to elucidate the correlation between the gene expression and IF, we looked for gene expression analysis and compared the TCGA data for breast cancer adenocarcinoma with lung adenocarcinoma using the GEPIA 2 database (http://gepia2.cancer-pku.cn/). To our surprise, for all the genes analyzed (SH3PXD2B, IL-6, CTTN, MMP2, and ADAM23), there was a consistent correlation between protein abundance and gene expression levels in different tissue types (Figure 5B). Therefore, these findings imply that the gene expression levels can be a reliable predictor of protein levels in these instances. Since the protein levels get significantly perturbed in normal and malignant states, they can be utilized as a potential biomarker for studying breast cancer-related malignancy.

### Bioluminescent IVIS imaging traces high metastatic migration in BCLM cells

In order to trace whether or not breast cancer cells metastasize to other distant organs, we successfully transfected breast cancer cells (MDA-MB-231) and BCLM cells (MDA-MB-468) with Addgene vector ID 119816, pLentipuro3/TO/V5-GW/EGFP-Firefly Luciferase vector (vector map is attached in Figure 6A), and the luciferase expression was visualized using D-luciferin substrate. To our surprise, BCLM MDA-MB-468luc2 substantially exhibited stronger bioluminescent signals than MDA-MB-231luc2 cells (Figure 6B). Following this, the bio-distribution analysis of the transfected cells was assessed in NOD-SCID mice. For the experiment, a control mouse was maintained, while two mice were injected with transfected MDA-MB-231luc 2 cells and MDA-MB-468luc2 cells via tail vein and mammary fat pad injection and imaged using the PerkinElmer IVIS Spectrum 2 optical imaging platform. As expected, the control mouse did not exhibit any bioluminescence. However, mice injected with transfected cells showed strong photon signals in the ventral view. Of which, mice with MDA-MB-231 luc2 cells exhibited localized and limited radius of dissemination, while mice with MDA-MB-468luc2 cells had a robust metastatic spread throughout the liver, lungs, and lymph nodes that corroborates our earlier findings on the high metastatic ability of MDA-MB-468 cells during malignant transformation (Figure 6C and supplementary data 6a).

**Figure 6:**
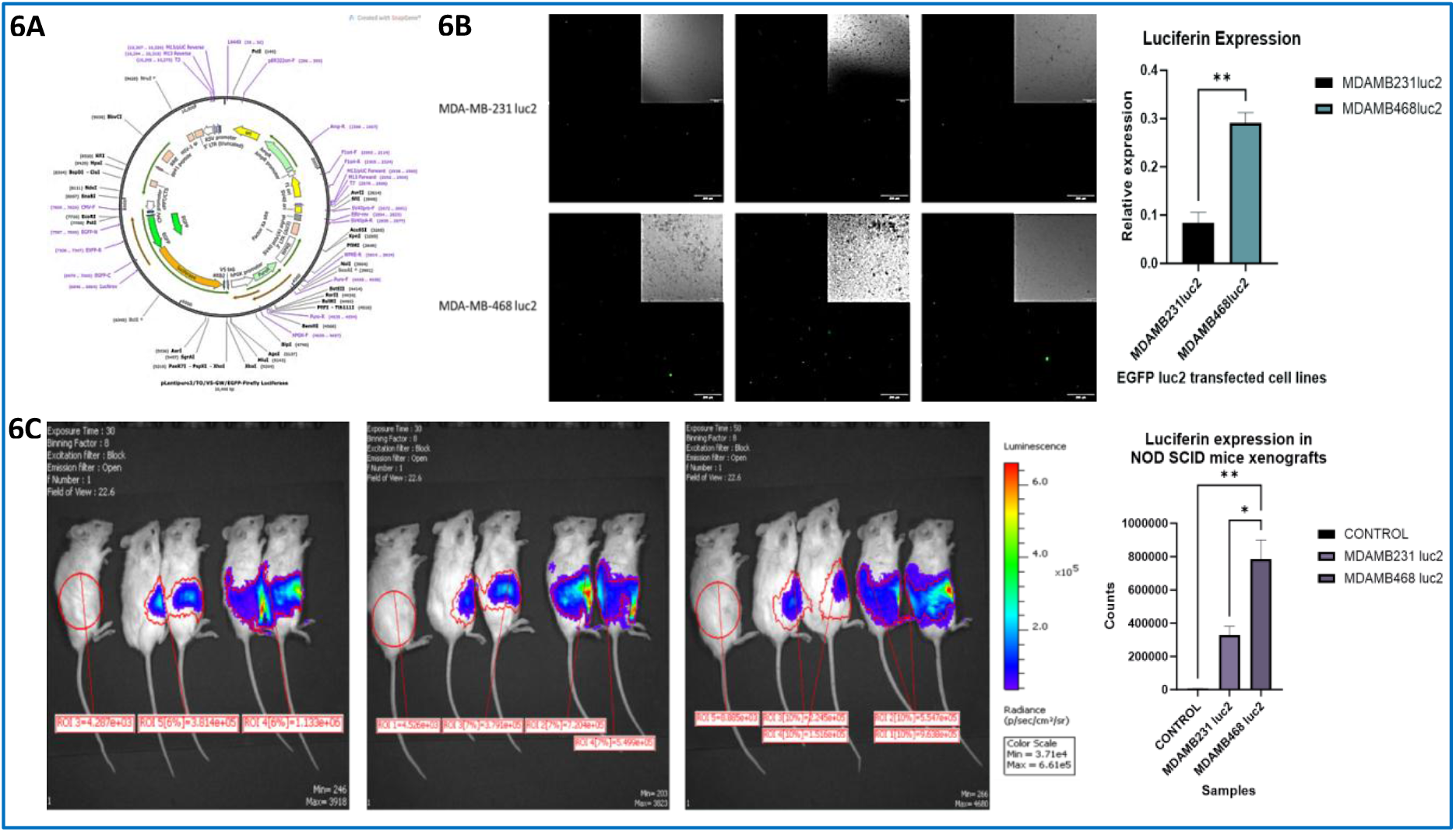
In vivo bioluminescence quantitation of MDA-MB-231 and MDA-MB-468 in mice model. (6A) A diagram illustrates a vector map that is utilized for transfection. (6B) represents successful transfection of metastatic breast cancer cells with a firefly luciferase vector. (6C) A figure represents the IVIS imaging of transfected MDA-MB-231 and MDA-MB-468 in experimental groups. In each experiment, mice with PBS (without cells) served as controls.

## Discussion

Metastasis continues to persist as a dreadful challenge in breast cancer patients, upholding its status as the primary cause of mortality among affected patients, despite significant advances in medical research and therapeutic strategies (21). Among the various forms of metastasis observed in breast cancer, the emergence of lung metastasis is alarming and potentially fatal. Current treatment modalities, including chemotherapy, radiotherapy, and surgical resection, are primarily palliative in nature, offering temporary relief rather than a permanent solution. This underscores the pressing need for innovative approaches to combat breast cancer lung metastasis effectively with a definitive cure (22). In recent years, the spotlight has shifted toward targeted therapies that are grounded in targeting key molecular players and signaling pathways underlying the intricate processes of metastasis, holding the potential to revolutionize our approach to treating BCLM.

The disruption of the proteostasis network is emerging as a key factor in breast cancer metastasis. Proteostasis refers to the maintenance of protein homeostasis, often through transcriptional and translational changes within cells. It involves the proper folding, assembly, trafficking, and degradation of proteins essential for preserving normal physiological conditions (23). Recent evidence suggests that disruption of the proteostasis network leads to many diseases including, cancer, at higher risk (24). It plays a crucial role in cancer cell survival and progression. Cancer cells experience proteotoxic stress due to the rapid synthesis of mutated or misfolded proteins, increased metabolic demands, and a hostile tumor microenvironment. To adapt and thrive, cancer cells often exploit the proteostasis network for their benefit (25). The disrupted proteostasis network and elevated proteotoxic stress may favor therapeutic resistance, invasion, and metastasis of breast cancer cells. Identifying key regulatory nodes and vulnerabilities in this network may lead to the development of novel therapeutic approaches for metastatic breast cancer.

SH3PXD2B plays a critical role in inducing EMT and metastatic processes in cancer cells (26). In this study, we embarked on comprehending the molecular landscape associated with breast cancer lung metastasis, with a particular focus on the role of SH3PXD2B as a potential key player in the process. The differential gene expression analysis unveiled a number of genes, including SH3PXD2B, and results indicated a contrasting expression of SH3PXD2B between BC and LM samples, which was further supported with the findings obtained from the mRNA expression data obtained from the CCLE database. The analysis also highlighted the prominence of SH3PXD2B expression, particularly in the MDA-MB-468 (TNBC) cell line, reinforcing its significance in metastatic carcinoma. Subsequently, the OS analysis indicated a poor prognosis for patients exhibiting increased expression of migratory markers in patients with BC and LC. The decline in the log-rank p-values below 0.05 indicates the significance of TNBC cells in cancer metastasis and its associated fatalities during specified time intervals. Additionally, the findings from TFTG and EMTome network analysis highlighted the role of SH3PXD2B’s interacting partners during BCLM. Consistent with the findings, the pathway enrichment analysis revealed key pathways regulated by SH3PXD2B and protein complex, validating their significant contribution to EMT and metastasis processes. Cytofunctional analysis shed light on the role of SH3PXD2B in influencing migratory events in breast and cancer cells, followed by a high expression of IL-6 in MBC patients, suggesting the additive role of these proteins in transformed tissues.

As SH3PXD2B has been the focus of our study and validated at multiple steps, we were curious to utilize the predicted secondary structures in order to employ an effective therapeutic strategy for MBC preventing BCLM. Since the explored structure predicted the highest level of confidence, targeting ordered regions in the protein structure having effective binding sites may enhance the biotherapeutics potential. As we elucidated SH3PXD2B as a master regulator of metastatic processes and ECM degradation, targeting SH3 domains could be a probable therapeutic target to assess patient-specific therapeutic interventions leading to personalized and effective therapies against BCLM.

## Abbreviations

ACTA2: ACTin Alpha 2
ADAM: A Distintegrin and Metalloproteinase
ALB: ALBumin
ARNT: Aryl Hydrocarbon Receptor Nuclear Translocator
BC: Breast Cancer
BCLM: Breast Cancer Lung Metastasis
BLK: B Lymphocyte Kinase
BRCA: BReast CAncer
CCLE: Cancer Cell Line Encyclopedia
CCNA2: CyCliN A2
CCSEA: Committee for Control and Supervision of Experiments on Animals
CDH: Cadherin
CDK2: Cyclin-Dependent Kinase 2
CDKN2A: Cyclin Dependent Kinase Inhibitor 2A
CEBPA: CCAAT/Enhancer Binding protein Alpha
CNOT3: CCR4-NOT transcription complex subunit 3
CPSF3: Cleavage and Polyadenylation Specificity Factor 3
CTNNB1: CaTeNiN Beta 1
CTTN: CorTacTiN
CYBA: CYtochrome B-245 Alpha chain
DMEM: Dulbecco’s modified Eagle’s medium
DUOX: DUal OXidases
EBF3: Early B-cell Factor 3
ECM: Extra Cellular Matrix
EGF: Epidermal Growth Factor
EGFR: Epidermal Growth Factor Receptor
EMT: Epithelial-Mesenchymal Transition
ESR1: EStrogen Receptor 1
FGR: Gardner-Rasheed Feline sarcoma
FN1: Fibronectin 1
FOX: Forkhead bOX
GATAD1: GATA zinc finger Domain containing 1
GFI1: Growth Factor Independent 1 transcriptional repressor
GRB: Growth factor Receptor Bound
IL: Interleukin
KLF1: Kruppel-Like Factor 1
L-15: Leibovitz’s-15
LC: Lung Cancer
LUAD: LUng ADenocarcinoma
MMP: Matrix Metalloproteinases
NFkB: Nuclear Factor kappa B
NOX5: NADPH Oxidase 5
NSCLC: Non-Small Cell Lung Cancer
PI3KCA: Phosphatidylinositol-4,5-bisphosphate 3-kinase catalytic subunit alpha
PrDOS: Protein Disorder Prediction System
PRPF: Pre-mRNA Processing Factor
RELA: v-rel avian reticuloendotheliosis viral oncogene homolog A
RPMI: Rosewell Park Memorial Institute
SH3PXD2B: SH3 and PX Domains 2B
ShScr: Short hairpin Scrambled
SNAI1: Snail family transcriptional repressor 1
SP1: Specific Protein 1
STAT3: Signal Transducer and Activator of Transcription 3
TCGA: The Cancer Genome Atlas
TFAP2A: Transcription Factor AP-2 alpha
TKS: Tyrosine Kinase Substrate
TNBC: Triple-Negative Breast Cancer
TPVG: Trypsin Phosphate Versene Glucose
VIM: VIMentin
ZEB: Zinc-finger E-box-Binding homeobox
ZNF589: Zinc Finger Protein 589
ZSCAN9: Zinc Finger and SCAN domain containing 9

