## Supplementary file 3 for "Unlocking the role of SH3PXD2B in epithelial-to-mesenchymal transition driving Breast Cancer Lung Metastasis"

Figure 1a

Supplementary data- 1

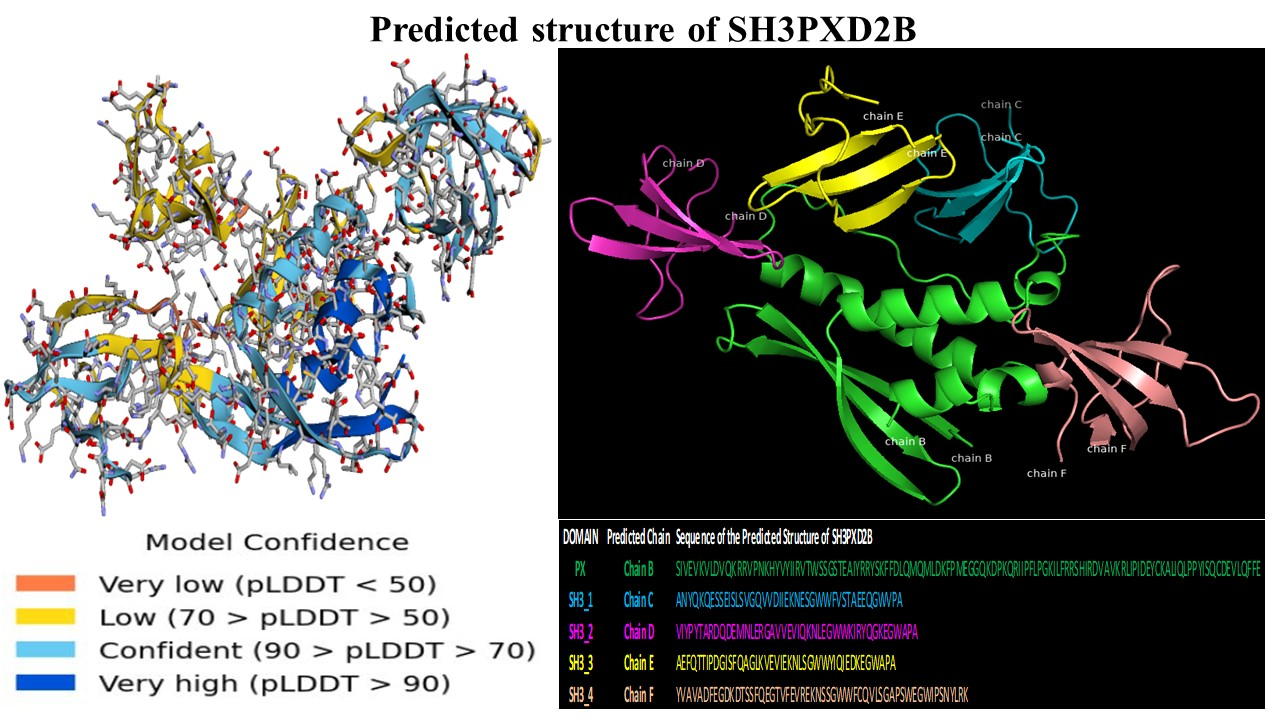

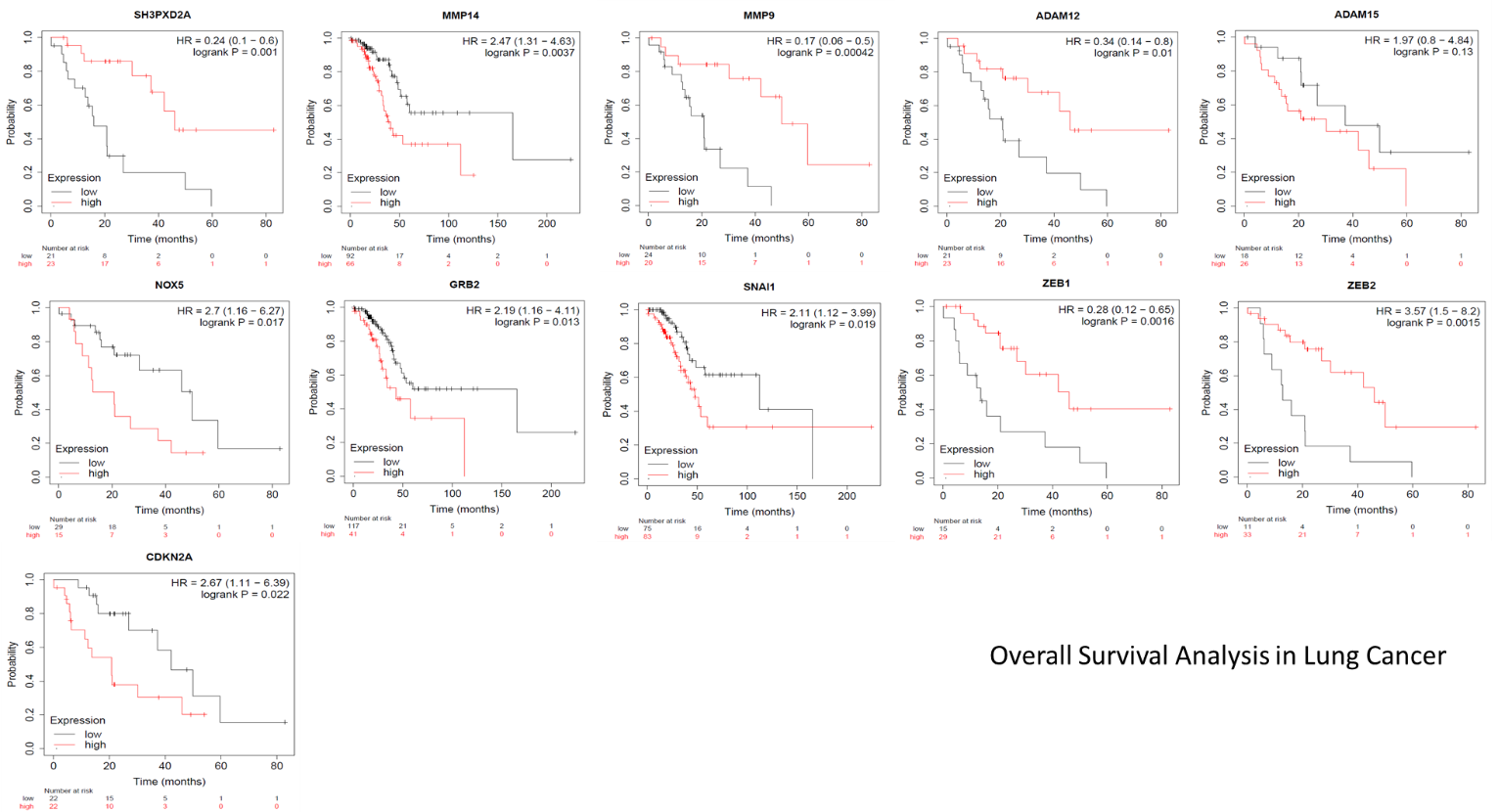

Supplementary data- 2

Figure 2a

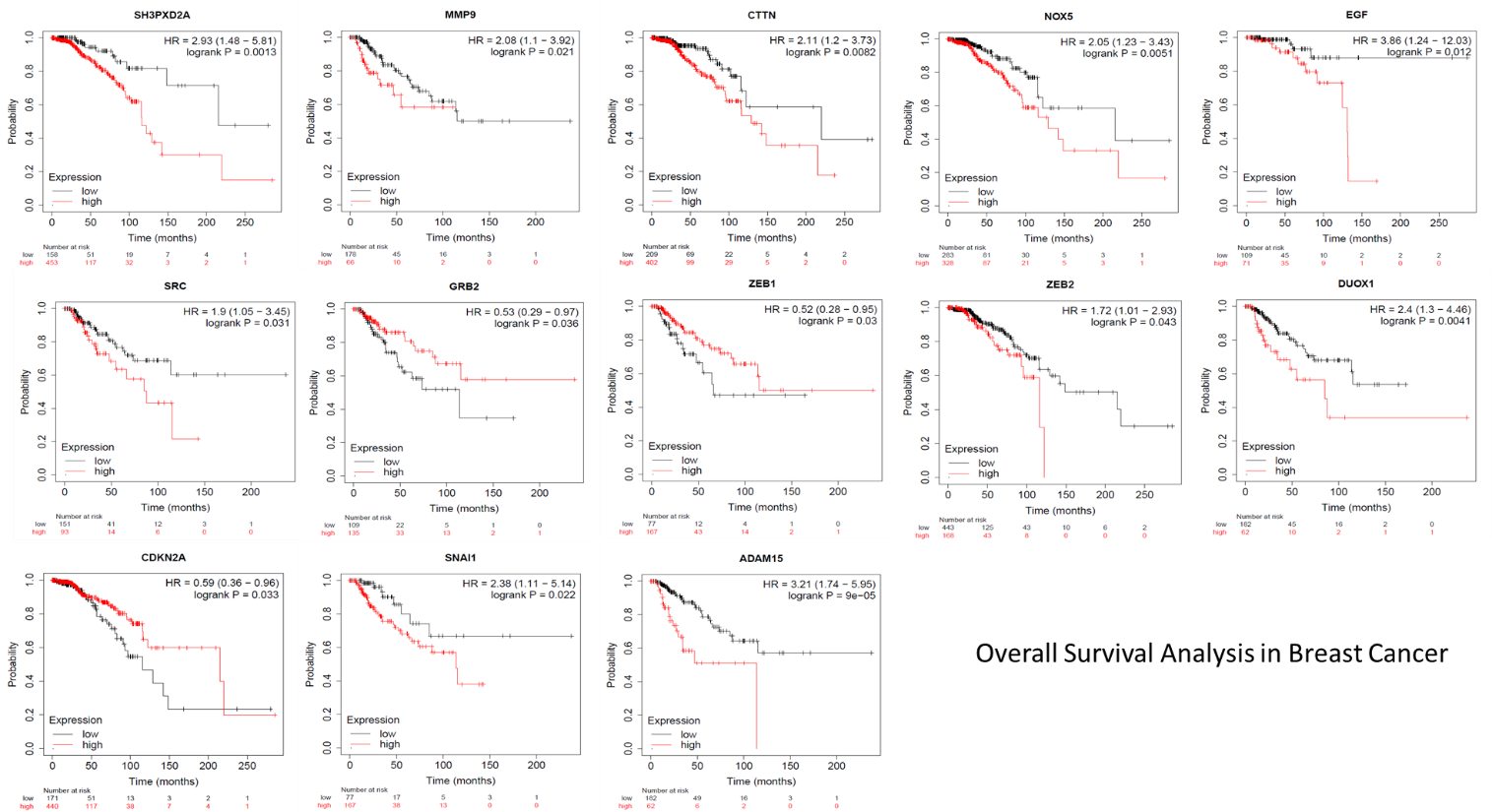

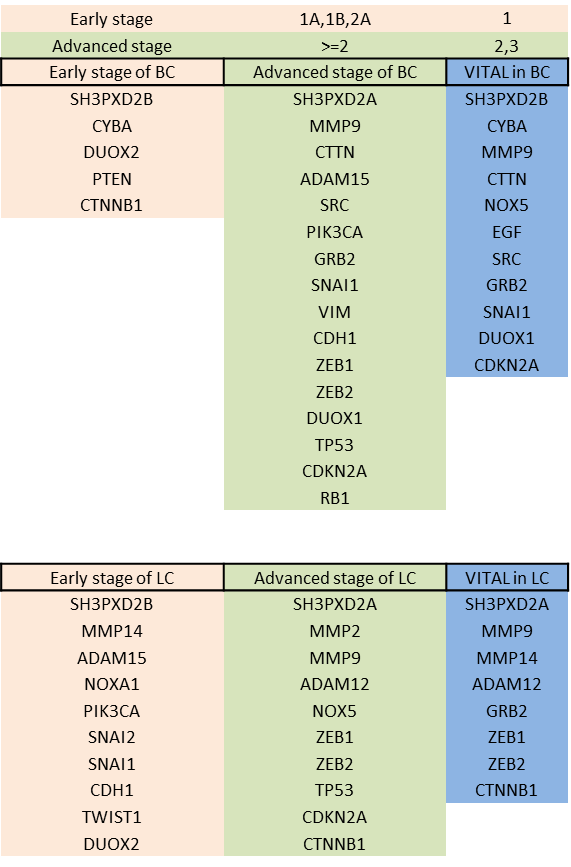

Figure 2b

Figure 2c

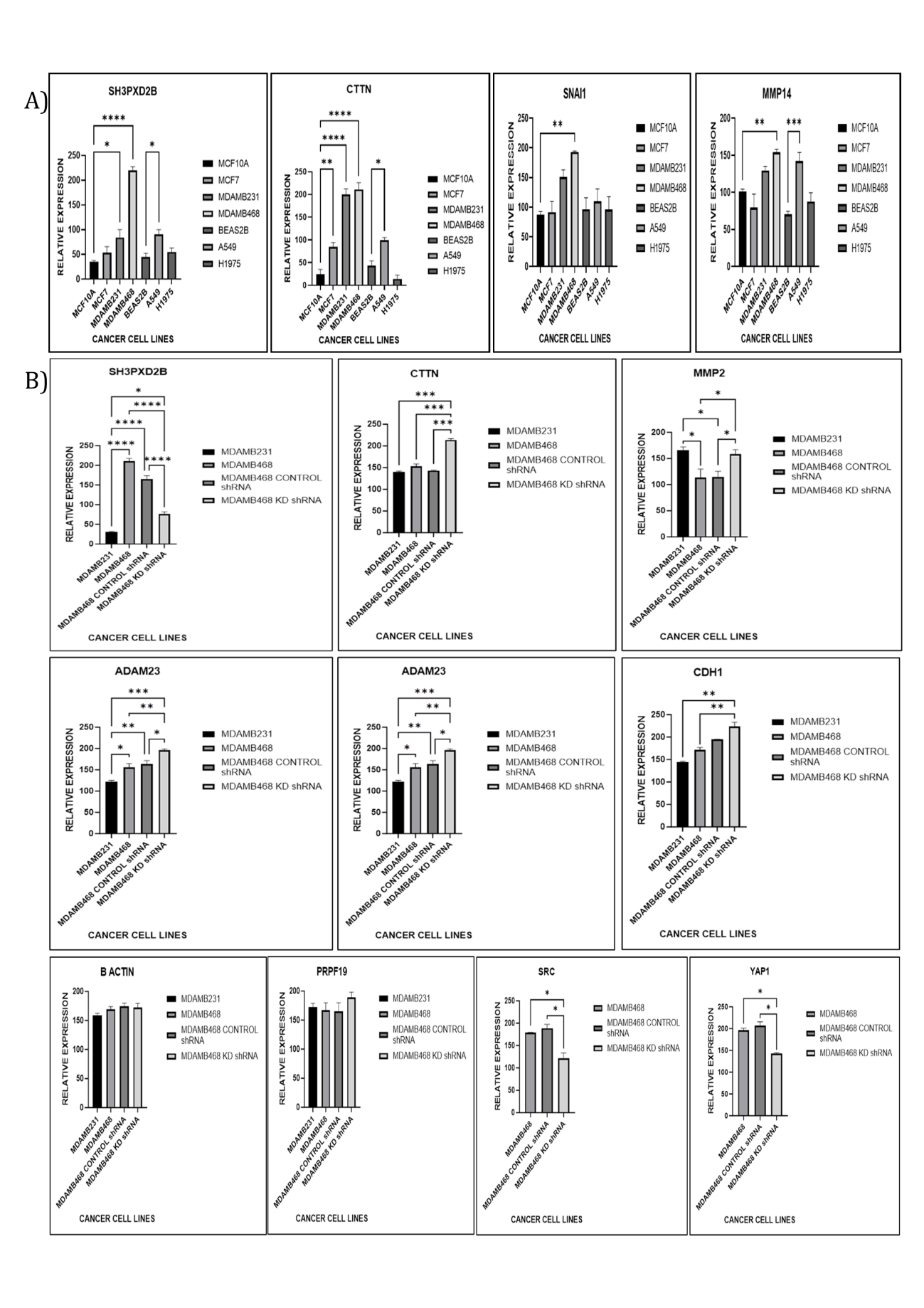

Figure 2d

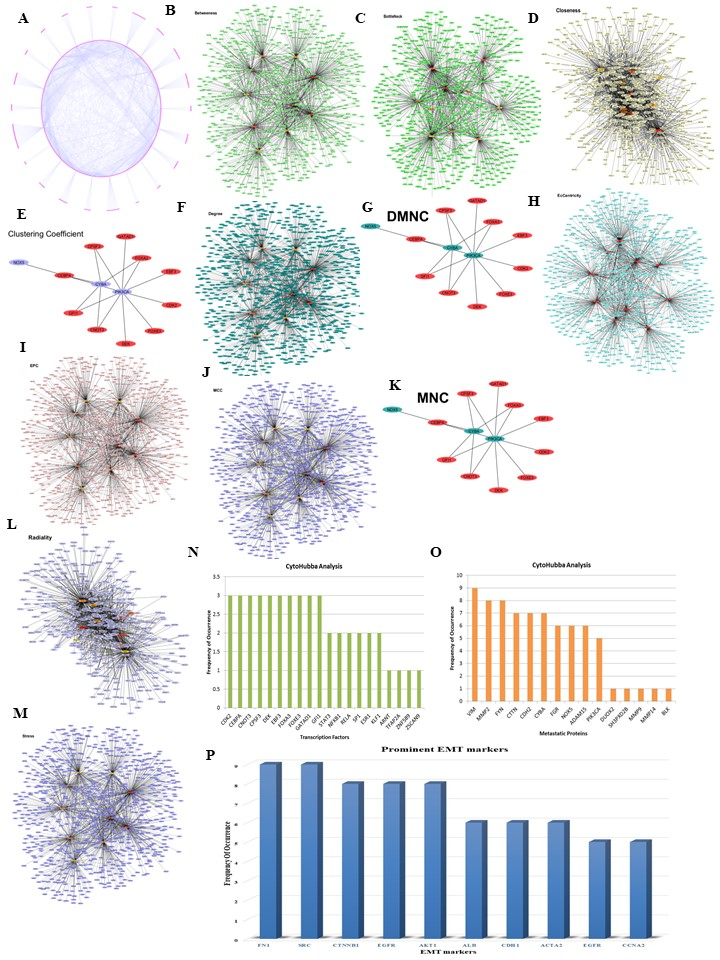

Separate excel sheet is provided

Data 3a

Supplementary data- 4

Supplementary data- 4

Figure 4a

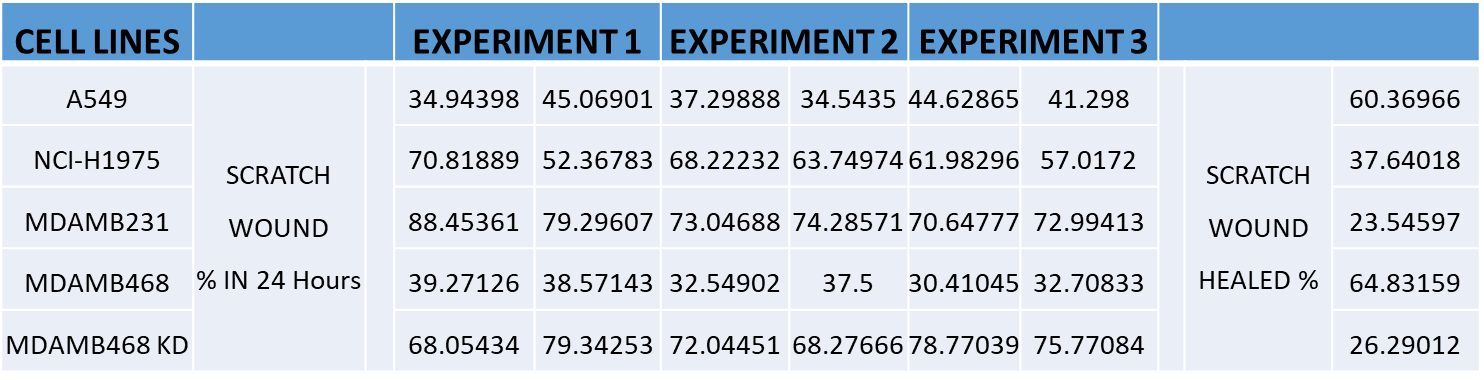

Figure 4b

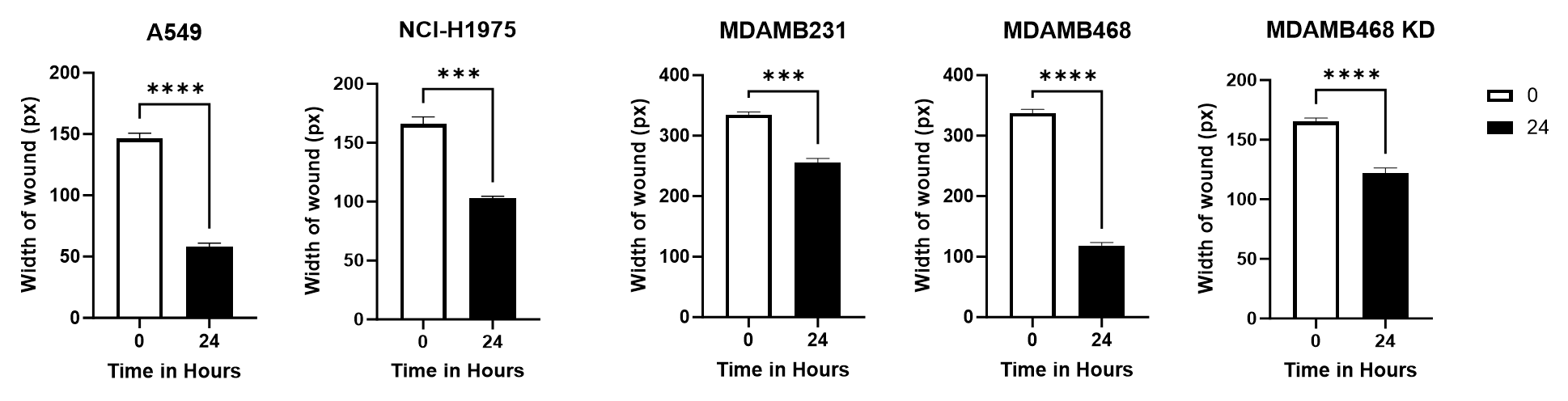

Figure 4c

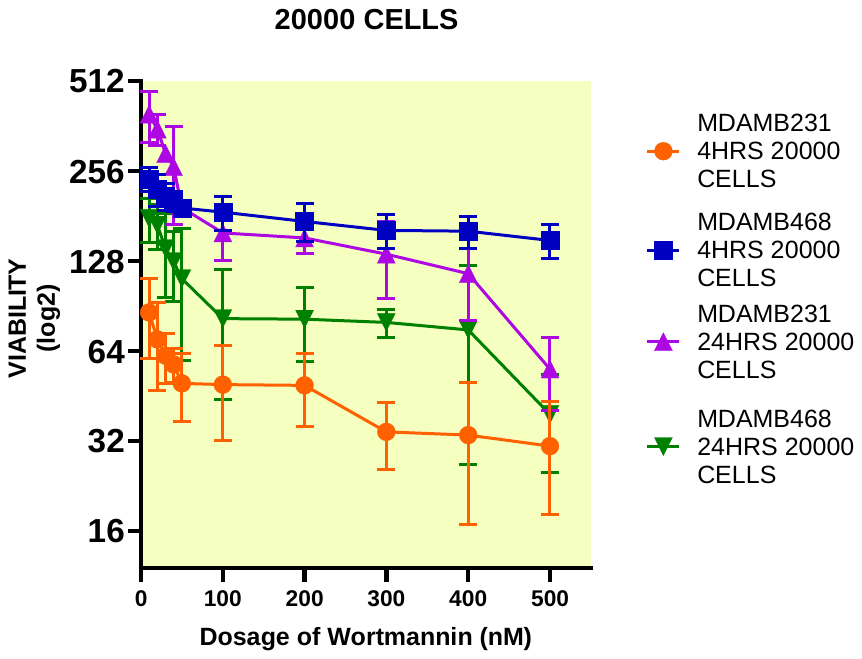

Figure 6a

Supplementary data- 6

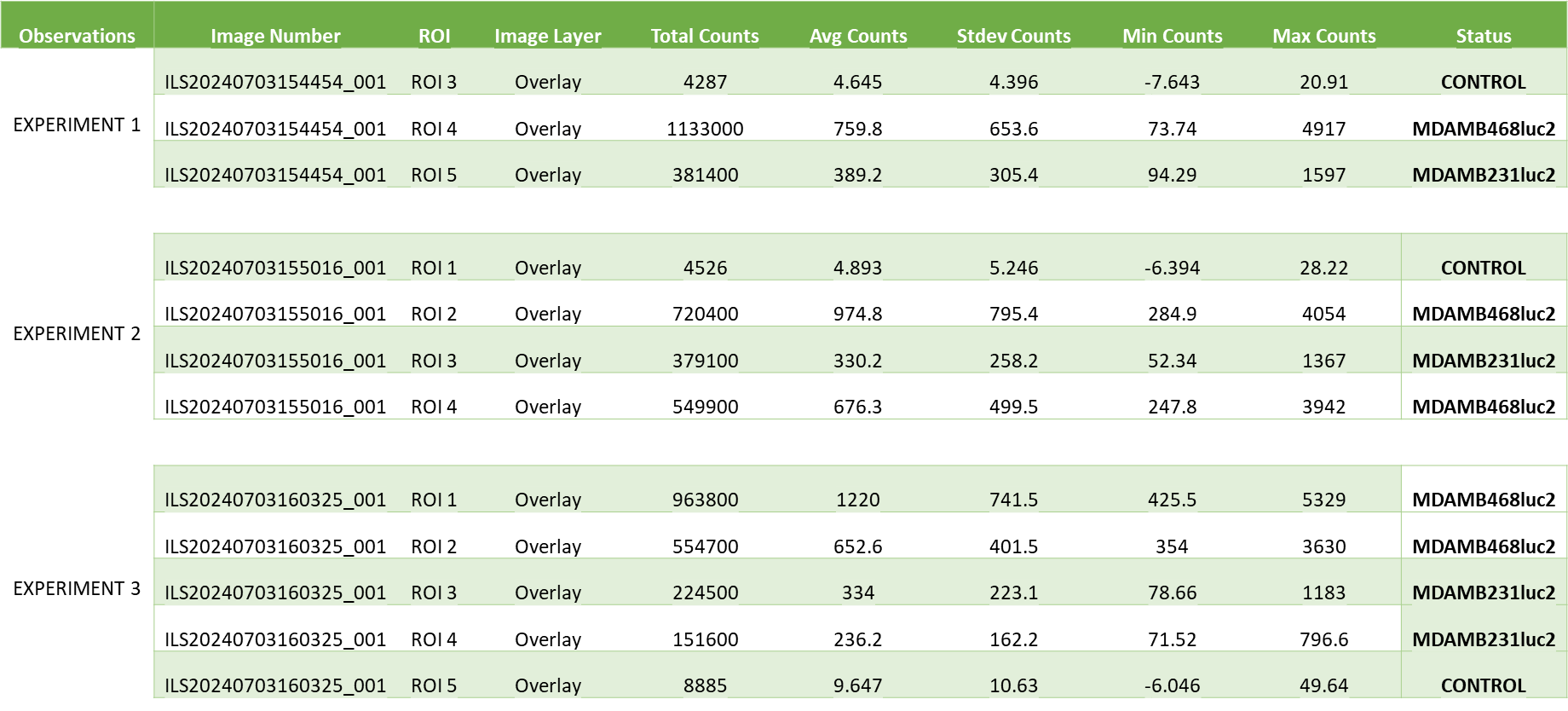
